# Innovative 3D-image analysis of cerebellar vascularization highlights angiogenic gene dysregulations in a murine model of apnea of prematurity

**DOI:** 10.1101/2025.10.17.683055

**Authors:** A. Rodriguez-Duboc, C. Racine, M. Basille-Dugay, D. Vaudry, B. Gonzalez, D. Burel

**Author notes:** Kavli Institute for Systems Neuroscience, Norwegian University of Science and Technology (NTNU), Trondheim, Norway. K. G. Jebsen Centre for Alzheimer’s Disease, Norwegian University of Science and Technology (NTNU), Trondheim, Norway. **Corresponding author: Pr Delphine Burel**.

## Abstract

Apnea of prematurity (AOP) affects 50% of preterm infants causing intermittent hypoxia (IH), which can lead to long-term neurodevelopmental deficits. Cerebellar abnormalities have been observed in AOP but the relationship between vascular alterations and neural development remains unclear. This study investigates how IH affects cerebellar angiogenesis using a murine model of AOP.

We developed an innovative 3D imaging workflow combining IMARIS and VesselVio software to quantitatively analyze cerebellar vascularization at different postnatal stages (P4, P8, P12, P21, and P70). We correlate these results with a transcriptomic analysis of 23 angiogenesis-related genes in the same stages to uncover the associated molecular pathways. We found that IH induced significant vascular changes, particularly at P4, with a global increase in vascular-network dimensions. By P8, the vascular network normalized, but genes were downregulated in all pathways studied. After P12, at the end of the IH protocol, transcriptional regulations vary but persist long-term. Moreover, differential analysis showed distinct effects on superficial versus deep vascular networks, allowing for a more precise understanding of remodeling patterns throughout development. Overall, transcriptomic changes were associated with morphological alterations in a time-dependent manner, suggesting a multiphasic IH response through development with lasting effects. Key regulations included VEGF, angiopoietin, and matrix metalloprotease signaling.

These findings demonstrate that IH disrupts cerebellar angiogenesis in parallel with neurogenesis, potentially contributing to the neurodevelopmental deficits observed in AOP. Thus, the interconnected nature of angio- and neurogenesis during cerebellar development makes it crucial to take vascular aspects into account in therapeutic approaches to neurodevelopmental disorders.

**Highlights:**

- Novel 3D imaging workflow reveals cerebellar vascular changes in apnea of prematurity mouse model
- Intermittent hypoxia induces early hypervascularization at P4 followed by later adaptation
- Superficial and deep vascular networks show distinct responses to intermittent hypoxia
- Vascularization-related genes are affected during cerebellar development
- Alterations in angiogenesis go together with neurodevelopmental defects

## Introduction

Apnea of prematurity (AOP) consists of an abnormal respiratory rhythm where breathing is interrupted at least every 5 minutes and for at least 20 seconds, thus inducing a state of intermittent hypoxia (IH). Premature newborns are particularly prone to AOP due to their immature respiratory system (Erikson et al., 2021; Moriette et al., 2010). This condition affects 50% of all preterm infants and nearly 100% of neonates born under 28 gestational weeks. The neonates will typically resume a normal breathing pattern by the corrected term, but several metadata provide evidence of a correlation between the duration of AOP and the incidence of developmental abnormalities that lead to long term behavioral deficits (Henderson-Smart, 1981; Janvier et al., 2004; Pergolizzi et al., 2022). Among the reported deficits, children suffer of motor coordination, language, and spatial orientation impairments, suggesting that a cerebellar alteration could be involved. According to this hypothesis, we have recently demonstrated, thanks to an established mouse model of AOP (Cai et al., 2012), that IH induces a delay of cerebellar cortex maturation and an alteration of the dendritic arborization of Purkinje cells, with long lasting transcriptional alterations (Leroux et al., 2022; Rodriguez-Duboc et al., 2023). The strong effect of AOP on the cerebellum is due to the developmental window of this nervous structure which mainly occurs after birth.

Indeed, in mice as in humans (Nieuwenhuys et al., 2008; Sillitoe and Joyner, 2007), cerebellar cortical development starts during embryogenesis from two germinative regions, the rhombic lip and the ventricular zone. Glutamatergic granule cells come from the rhombic lip and migrate tangentially to form the transitory external granular layer (EGL) at the surface of the cerebellum. GABAergic Purkinje cells (and several types of interneurons) arise from the ventricular zone and migrate transversally to the surface to create a monolayer of Purkinje cells just below the EGL. As a result, at birth, the cerebellum is composed of 3 layers, the EGL, containing granule cell precursors (GCP), and the Purkinje cell layer (PCL) separated by the molecular layer (ML). However, at this point the structure is still immature and has yet to undergo postnatal maturation. Firstly, the GCP proliferate intensively and cross the ML and the PCL to form the definitive granule cell layer (GCL) until the total disappearance of the EGL. Meanwhile, the GCP finish their maturation and send their axons, called parallel fibers, to the ML. At the same time, Purkinje cells develop their dendritic trees into the ML to contact parallel fibers. Much like in the rest of the brain, the neurohistogenetic process is mirrored by angiogenesis within the cerebellar cortex. Then cerebellar vascularization initiates in the perineural vascular plexus that covers the neural tube (Vogenstahl et al., 2022). Then, collaterals from pial vessels enter the cerebellar cortex and irrigate the EGL. At first sparse in the EGL during the proliferative phase of GCP, blood vessels then continue branching out and the number of capillaries increases in the ML and the GCL (Mecha et al., 2010). Therefore, as hypoxia is known to remodel the vascular organization and enhance the density of blood vessels in the brain (Guan et al., 2022; LaManna et al., 2004), we can hypothesize that AOP could disrupt vascularization during the postnatal development of the cerebellum (as it does with neuronal cells).

In line with this hypothesis, our previous transcriptomic results, showed that the expression of the hypoxia inducible factor 1 alpha (HIF1α) is affected in response to our IH protocol (Leroux et al., 2022). HIF1α is considered as the main regulator of angiogenesis since this factor responds to low levels of dioxygen (O_2_) by promoting the expression of pro-angiogenic actors, including the vascular endothelial growth factor (VEGF) and platelet derived growth factor β (PDGFβ) as well as miRNA (Monaci et al., 2024). Another angiogenic process could also occur in the brain via the Ang2/Tie2 pathway. Indeed, under hypoxic conditions, endothelial cells increase Ang2 production which acts in an autocrine manner on Tie2 receptor to destabilize capillaries and promote angiogenesis (Augustin et al., 2009). Interestingly, Tie2 has been recently described in Purkinje cells and it has been demonstrated that the neuro-vascular signalling Ang2/Tie2 controls dendritic morphogenesis of Purkinje cells during cerebellar postnatal development (Luck et al., 2021).

All these data led us to consider that the alteration of the Purkinje dendritic trees observed after IH in our AOP model could be linked to a perturbation of cerebellar vascularization. To provide elements in understanding this mechanism, we performed a transcriptomic study focused on the main factors involved in cerebellar angiogenesis at different postnatal ages. Moreover, to correlate putative modification of gene expression with defaults in vessel morphology, we developed an image analysis workflow using both IMARIS and VesselVio software to visualize the cerebellar vascularization in 3D during postnatal development and obtain comparable quantitative parameters between normoxic and IH conditions.

## Materials and methods

### Animals

Animals used in this study were wild type C57Bl6/J mice born and bred in an accredited animal facility (approval number A 76-451-05). Animal experiments were approved by the French Regional Ethics Committees and the Ministry of Education, Research and Innovation (project n°10310-2017041811326427v5 – 13/02/19 - 12/02/22), and they were performed in accordance with the European Committee Council Directive (2010-63-EU). Mice lived on a 12-hour light/dark cycle and had free access to food and water. Sex was determined based on anogenital distance measurement as well as pigment-spot localization (Wolterink-Donselaar et al., 2009). From the postnatal day 2 (P2) onwards, mice were assigned a unique identifier before initializing the IH protocol. There was no blinding in this study, and sample size was chosen based on power determination from our preliminary studies (Leroux et al., 2022; Rodriguez-Duboc et al., 2023). A total of 28 litters of animals including the dams were used for this study.

### Intermittent hypoxia protocol

Our IH protocol relies on a custom hypoxia chamber, which is based on the protocol developed by Cai et al., and has been validated to mimic AOP (Cai et al., 2012; Leroux et al., 2022; Rodriguez-Duboc et al., 2023). IH was achieved by the repeated succession of 2-min cycles of hypoxia (5% O_2_; 20 s/cycle) and reoxygenation, for 6 hours per day, while mice were in their sleep phase (10 am - 4 pm). The protocol was initiated on neonatal P2 pups (IH group) until the desired stage, or for 10 days maximum. Throughout the protocol, the following parameters were constantly monitored in the chamber: oxygen concentration, hygrometry, temperature, and atmospheric pressure. The control/normoxia group (N) was placed in a chamber open to room air but mimicking the hypoxia chamber’s environment to control for external stressors.

### Sample gathering

For the transcriptomic study, mice were sacrificed at stages P4, P8, P12, P21, and P70 by decapitation after anesthesia by isoflurane inhalation (Iso-VET). Whole brains were immediately harvested and set in pure isopentane at -30°C. They were then stored in sterile containers at -80°C until further use. Sample sizes for RT-qPCR were: 13 for P4, of which 7N (3F+4M) and 6IH (4F+2M); 16 for P8, of which 6N (3F+3M) and 10IH (6F+4M); 25 for P12, of which 15N (8F+7M) and 10IH (2F+8M); 17 for P21, of which 9N (3F+6M) and 8IH (5F+3M); and 8 for P70, of which 5N (1F+4M) and 3IH (1F+3M).

For imaging studies, brains from P4 mice were directly harvested and fixed by immersion in 4% paraformaldehyde, given their small size. For P8, P12, P21, P70 stages, mice were lethally anesthetized by intraperitoneal injection of ketamine (100 mg/kg) and xylazine (10 mg/kg), and then sacrificed by intracardiac perfusion of NaCl 9‰ and paraformaldehyde 4%, before removing the brains. Brains were then postfixed overnight in 4% paraformaldehyde, and stored in phosphate buffer saline (PBS). Sample sizes for the clearing experiments were: 8 for P4, of which 4N (2F+2M) and 4IH (2F+2M); 8 for P8, of which 4N (2F+2M) and 4IH (1F+3M); 7 for P12, of which 3N (1F+2M) and 4IH (1F+3M); and 7 for P21, of which 3N (2F+1M) and 4IH (0F+ 4M).

### Panels and primer design

The vascularization gene panel was built based on an analysis conducted with the Cytoscape software (v.3.9.1; Shannon et al., 2003) using protein interaction data retrieved from STRING-DB (Szklarczyk et al., 2021). The panel was further expanded by enrichment with the plugin stringApp (v 2.0.1; settings: maximum additional interactors = 10, confidence cutoff = 0.40; Doncheva et al., 2019) and then cross referenced with ClueGO pathway (v.2.5.9; Bindea et al., 2009) and functional data retrieved from the literature. For readability and ease of interpretation, these findings are summarized in Table 1, and the resulting functions were used to group the quantitative RT-PCR results.

**Table 1.** Selected angiogenesis-associated genes, the functions they are associated with, and their roles within these functions. Angpt1: angiopoietin 1; Angpt2: angiopoietin 2; Anpep: alanyl aminopeptidase; Cdh5: vascular epithelium-cadherin; Col1a1: collagen, type I, alpha 1; ECM: extracellular matrix; F3: coagulation factor III; Fgf2: fibroblast growth factor 2; Flk1: vascular endothelial growth factor receptor 2; Flt1: vascular endothelial growth factor receptor 1; Lect1: chondromodulin; Mmp2: matrix metallopeptidase 2; Mmp9: matrix metallopeptidase 9; Ndp: Norrie disease protein; Nrp1: neuropilin 1; Pgf: placental growth factor; Serpine1: plasminogen activator inhibitor 1; Serpinf1: pigment epithelium-derived factor; Tek: endothelial-specific receptor tyrosine kinase; Tgfb1: transforming growth factor, beta 1; Thbs1: thrombospondin 1; Tie1: tyrosine kinase with immunoglobulin-like and EGF-like domains 1; Timp1: tissue inhibitor of metalloproteinase 1; Vegfa: vascular endothelial growth factor A.

| **Function abbreviation** | **Functional pathway** | **Genes with a putative positive effect on the function** | **Genes with a putative negative effect on the function** |
| --- | --- | --- | --- |
| VSM | Vascular stabilization and maturation | Tgfb1; Flk1; Angpt1; Tek; Vegfa | Tie1 |
| EEP | ECM and endothelial permeability | Cdh5; Vegfa; Mmp2; Mmp9; Angpt2 | Angpt1 |
| VIS | Vascular invasion and sprouting | Cdh5; Fgf2; Vegfa; Serpine1; Nrp1; Anpep; Mmp9; Ndp; F3 | Flt1; Col1a1; Thbs1; Timp1 |
| EPS | Endothelial proliferation and survival | Pgf; Tgfb1; Flk1; Nrp1; Angpt1; Tie1 | Lect1; Angpt2; Thbs1; Serpinf1 |
| VRP | Vascular remodelling and patterning | Flt1; Tie1 | Vegfa |
| HIA | Hypoxia-induced angiogenesis | Mmp2; Mmp9; Tek; Tie1; Vegfa; Serpine1 | Serpinf1 |

Gene primers were designed with the Primer Express software (v3.0.1; ThermoFisher Scientific) using nucleotide sequences from the NCBI Pubmed database. Primer pairs were ordered from Integrated DNA Technologies and their specificity was validated by linear regression of serial dilution data. Primer pairs were chosen preferentially to be on exon joining sites, with the least possible hairpin and dimer formation, and with similar size, GC percentage, and melting temperature for forward and reverse primers. Each sequence was blasted on Basic Local Alignment Search Tool (Blast, Ye et al., 2012) to ensure specificity. See Supplementary tables 1 and 2.

### RNA Extraction

The samples were homogenized in 1 ml of Trizol (ThermoFisher) and mRNAs were purified on column using the Nucleospin RNA extract II from Macherey-Nagel (cat. 740 955 250) according to manufacturer recommendations. RNA quantity and purity were analyzed by UV spectrophotometry (Nanodrop Technologies). The optical density (OD) of RNA was read at 230, 260, and 280 nm. The ratios OD 260 nm/OD 280 nm and OD 260 nm/DO 230 nm were calculated as indicators of protein, and salt/ethanol contamination, respectively. Ratio values between 1.6 and 2.0 were considered acceptable. As per the MIQE guidelines (Bustin et al., 2009), mRNA quality assessment was performed by a bioanalyser gel electrophoresis on RNA 6000 Pico chips (cat. 5067-1513, Agilent), and samples with an RNA integrity number (RIN) between 7 and 10 were considered qualitative enough to be analyzed. The mRNAs were then stored at -80°C until use.

### Quantitative RT-PCR

Messenger RNAs were converted to cDNA by reverse transcription using the Prime Script RT reagent kit (cat. RR037A, Takara). Relative gene expression level determination was done by real time PCR in 384-well plates. Total reaction volume was 5 μL including: 2.5 μL 2X Fast SYBR Green PCR Mastermix (cat. 4385612, Thermofisher), target gene-specific sense and antisense primers (0.15 µl of each, 100 nM final concentration), 1μL PCR-grade water, and 1.2 μL of sample solution. The cDNA samples and reaction mixes were distributed via the Bravo 1 liquid handling platform (Agilent). The real time PCR reaction took place in a QuantStudio Flex 12k thermal cycler (Applied Biosystems). For each gene of interest (GOI), sample measurements were conducted at least in duplicate and with a minimum of two housekeeping genes (HKG). Values were calculated via the 2^(-ΔΔCq)^ method where:

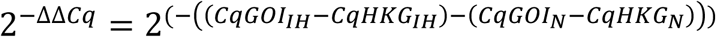

### Imaging

Cerebella previously fixed with 4% paraformaldehyde underwent a clearing protocol consisting of the following steps. Samples were dehydrated by being submerged in increasing concentrations of methanol (MeOH; 20%, 40%, 60%, 80% and 100%). Dehydrated samples underwent bleaching in a solution of 5% H_2_O_2_ (hydrogen peroxide) and 95% MeOH for 24 hours in order to decrease tissue autofluorescence. Samples were then rehydrated by being submerged in decreasing concentrations of MeOH (80%, 60%, 40% and 20%), then washed in solution PTx.2 (100 mL PBS 10X + 2 mL Triton-X100 + Q.S. distilled water). Samples were then permeabilized for one day at 37 °C under agitation with a solution containing 1X PBS, 0.2% TritonX-100, 20% dimethyl sulfoxide (DMSO), glycine (23 mg/mL) and thimerosal at 0.1 g/L (antifungal). Finally, non-specific binding sites were blocked with a 1X PBS solution containing 0.2% Triton-X100, 10% DMSO, 6% normal donkey serum (NDS) and thimerosal for one day at 37 °C under agitation.

Thereafter, the brains were incubated with the primary antibodies diluted in PTwH solution (100 mL PBS 10X + 200 μL Heparin (50 mg/mL) + Q.S. distilled water), containing 5% DMSO and 3% NDS at 37 °C under agitation for 6 days. After 6 rinses with PTwH at room temperature, the samples were incubated for 5 days with the appropriate secondary antibodies (Table 2) diluted in PTwH containing 3% NDS at 37°C under agitation. The brains were then rinsed several times with PTwH at room temperature under agitation, followed by dehydration in MeOH baths of increasing concentration (20%, 40%, 60%, 80% and 100%). A delipidation of the brains was then performed by incubation in a solution containing 66% dichloromethane (DCM) and 33% MeOH for one night under agitation and then in DCM 100% for 30 minutes. These last two steps homogenize the refractive indexes of the cellular structures and induce their transparency once placed in dibenzylether.

**Table 2.** Antibodies used for the visualization of blood vessels. DAG: donkey anti-goat; DARt: donkey anti-rat; α-SMA: smooth muscle actin alpha; PECAM1: platelet endothelial cell adhesion molecule 1. The secondary antibodies were purchased from Molecular Probes.

| Primary Antibodies | Target | Dilution | Species | Supplier | Secondary antibodies | Dilution |
| --- | --- | --- | --- | --- | --- | --- |
| Podocalyxin | Capillaries | 1:200 | Goat | R&D Systems (#AF1556) | DAG-Alexa 594 | 1:400 |
| α-SMA-Cy3 | Arteries | 1:500 | Mouse | Sigma-Aldrich (#C6198) | N/A | N/A |
| PECAM1 | Arterioles | 1:200 | Rat | Millipore (#CBL1337) | DARt-Alexa 594 | 1:400 |

The 3D acquisitions of the transparent cerebella were performed on the Ultramicroscope Blaze (Miltenyi Biotec) using the software ImspectorPro. Image analysis and vessel modelling were done using the Imaris software (Oxford Instruments, version 10.0) and then transferred to VesselVio 1.2 software to obtain quantitative and comparative parameters (Bumgarner et al., 2022).

### Statistical analysis

Statistical analyses were performed within the R statistical computing environment (version 4.3). Both real time PCR and imaging data were modeled through the Generalized Linear Mixed Model (GLMM) framework, using the {glmmTMB} package (Brooks et al., 2017). For real time PCR data, a Gaussian likelihood with an identity link function was used to model the distribution of the DCq samples for each gene of interest. When DCq samples for one gene were split over multiple plates, a random intercept was added to account for intra-plate correlations.

Model diagnostics were done using the {DHARMa} (Hartig, 2022) and {performance} (Lüdecke et al., 2021) packages. The fitness of each model was assessed through both visual checks (e.g., posterior predictive checks, QQ plots, and residuals vs predicted values) and quantitative indices of model fit (e.g., AIC: Aikake Information Criterion). When several competing models were possible a priori, we selected the most plausible one primarily based on our theoretical understanding of the response’s properties, and, to a lesser extent, to minimize AIC and favor model parsimony.

Contrasts and p-values for relevant hypotheses were obtained using the {emmeans} package (Lenth, 2022). They were computed on the link scale, using Wald t-tests, without any multiplicity adjustments. For all analyses, p < 0.05 was considered significant and for each figure asterisks indicate the level of statistical significance: one for p < 0.05, two for p < 0.01, and three for p < 0.001.

## Results

### Workflow of the cerebellar vascularization analysis

Thanks to a concomitant triple labelling (smooth muscle actin alpha (α-SMA), platelet endothelial cell adhesion molecule 1 (PECAM1) and podocalyxin), we were able to visualize all the vascular networks, namely, the arteries, veins, and capillaries, at all stages of postnatal development (Figure 1). The labelling is present in all cerebellar structures. Signal intensity is stronger in the cortex than in the depth of the tissue at P12 and P21 because of the higher volume of the sample that impedes antibody penetration despite applying an increasing permeabilization protocol. Due to the variability of the cerebellum’s morphology, no method currently exists to analyze the vasculature of postnatal mice cerebella. Therefore, we developed the following workflow by optimizing each step of analysis using Imaris and Vesselvio software.

**Figure 1:**
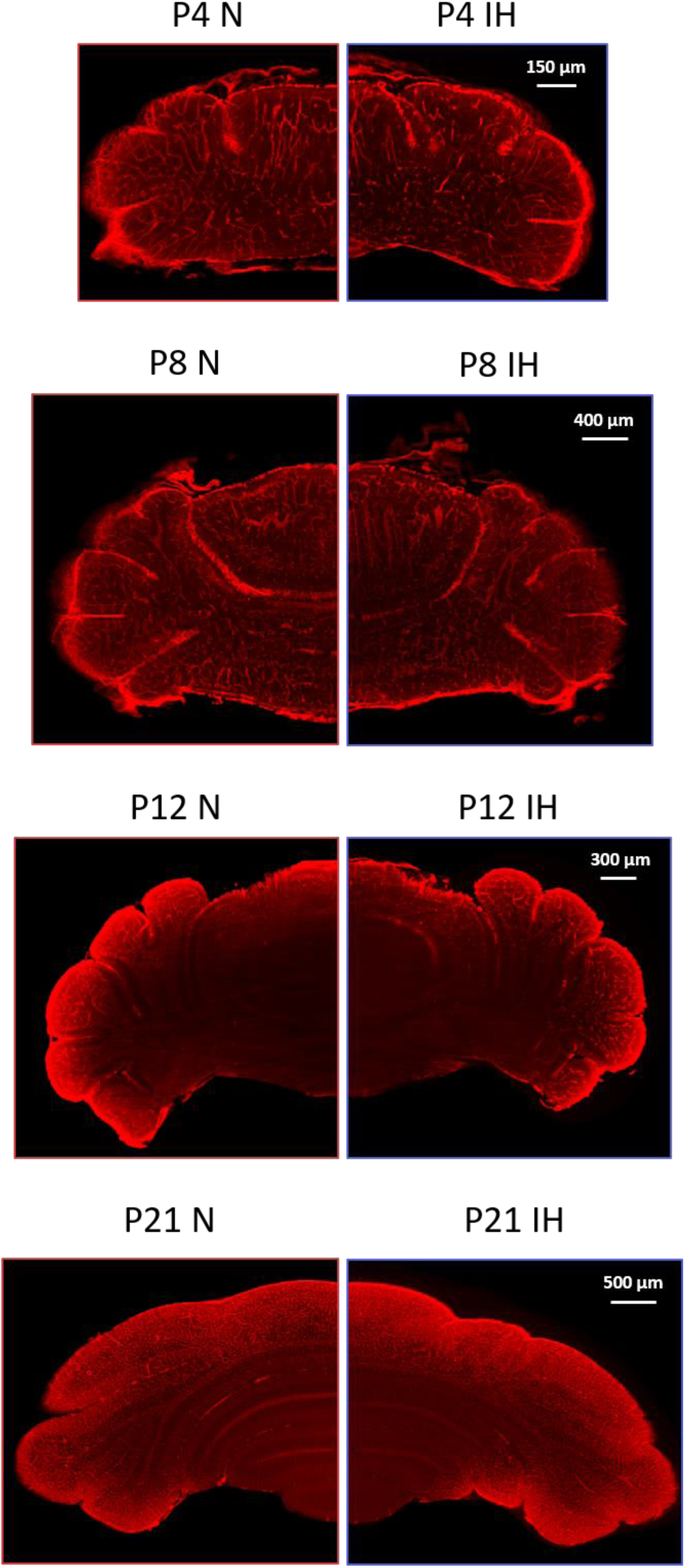
Images illustrating the cerebellar vascularization in normoxia and intermittent hypoxia conditions. Lightsheet microscopy 2D images of cerebella illustrating the vascularization in control (N) and hypoxic mice (IH) at various postnatal stages (P4, P8, P12, and P21). Whole cerebella were simultaneously incubated with antibodies against PECAM, podocalyxin and a-SMA before clearing and image acquisition. For each stage, the scale is indicated in the top right corner. IH: intermittent hypoxia; N: normoxia; Px: postnatal day x.

The file resulting from the 3D lightsheet acquisition was converted to an *.ims* format, then the *Surface* function of the Imaris software was used to delineate de cerebellum from the whole hindbrain 3D acquisition (Figure 2A). Because of the complexity of the cerebellum, the delineation was performed manually, by drawing cerebellar outlines every 20 µm in the 2D-slices mode (Figure 2B). The resulting *Surface* was used to obtain a channel (mask) including only the cerebellum (Figure 2C), which allows the calculation of the total cerebellar volume as well as the total volume of the vasculature as follows. The vascularization modelling was built with the *Filament* modality and applied to the cerebellum channel. *The Automated Autopath Algorithm* mode was chosen among the six algorithms proposed by the software because this process models dense networks with numerous branches. The algorithm is comprised of three steps and is primed on a representative region of interest.

**Figure 2:**
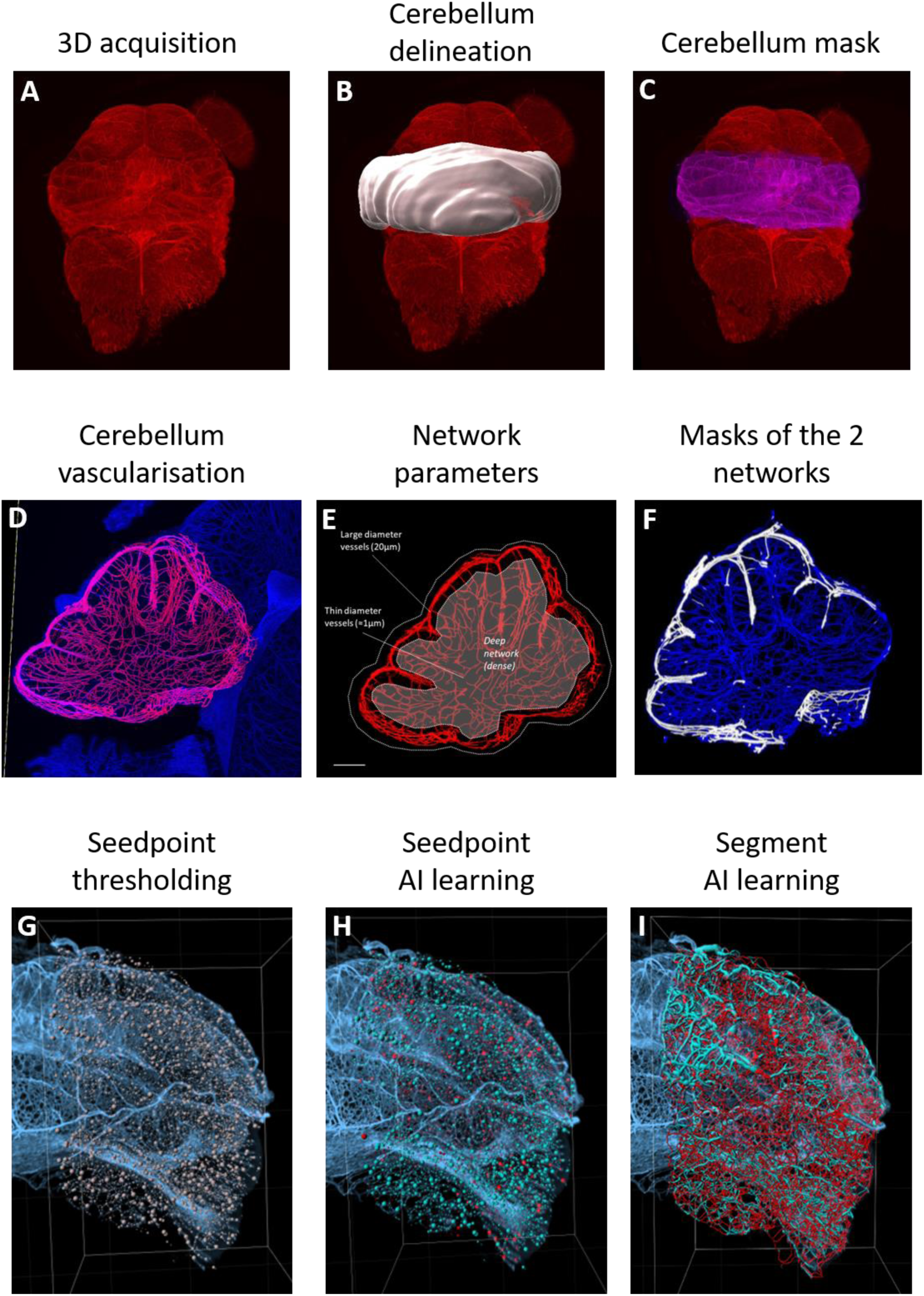
Images illustrating the IMARIS workflow developed for the vascular network modeling. Sequential workflow steps allowing the analysis of the cerebellar vascular network of a P4 mouse cerebellum on the Imaris software. From a 3D lightsheet acquisition (A), the cerebellum is delineated (B) and a mask is created (C). Within that selected volume, the cerebellar vascularization is segmented (D), which allows the network visualization (E) and the separation of a deep and a superficial network (F). Then the threshold of seedpoints is defined (G), and thanks to the artificial intelligence module (AI), Imaris is able to discriminate “true” (blue) and false (red) seedpoints (H), and “true” (blue) and false (red) segments (I). Px: postnatal day x.

The first step involves choosing the detection modalities. The observation of the vascular organization on a sagittal cerebellar section revealed a high heterogeneity of vessel diameter (Figure 2D). Large vessels are located on the surface of the cortex, and fine vessels are in the center of the cerebellum (Figure 2E). Thus, to optimize the quality of the modelling beforehand, we split the vascularization into two networks based on a threshold of voxel intensity. All the positive voxels were considered as the “superficial network” and all the negative voxels represented the “deep network”, which includes all the vessels in the organ’s core (Figure 2F). The second step of the protocol involves defining the diameter range to be considered. These diameters constitute the detection threshold for branching points, defined as the junctions between each vascular segment. The network splitting allowed more homogeneity of the vessel diameters. However, we chose a *Multiscale* option to guarantee the detection of all vessels. Thus, a diameter range of [1 - 10µm] was chosen for the “deep network” and a [1 - 20µm] range for the “superficial network.”

The third step of the algorithm requires training the artificial intelligence (AI) in three different phases. In the first phase, the aim is to define the threshold for detecting orientation changes. This step is crucial because it enables the subsequent training of the software to classify these points and then generate the segments. So the wider the range is, the more efficient the training will be (Figure 2G). In the second phase, based on the threshold, the AI predicts keeping or discarding seedpoints. The AI can be further trained by correcting potential choice errors until a satisfactory result is achieved, but the correction must not exceed 100 manual points to avoid conflicting information, which could disrupt the training (Figure 2H). With the same method, the third phase consists of classifying and selecting the resulting segments (Figure 2I). Finally, the process is applied to the entire 3D volume (Figure 3A). For each postnatal stage, the same AI parameters were saved and re-used on all normoxic and hypoxic cerebella.

**Figure 3:**
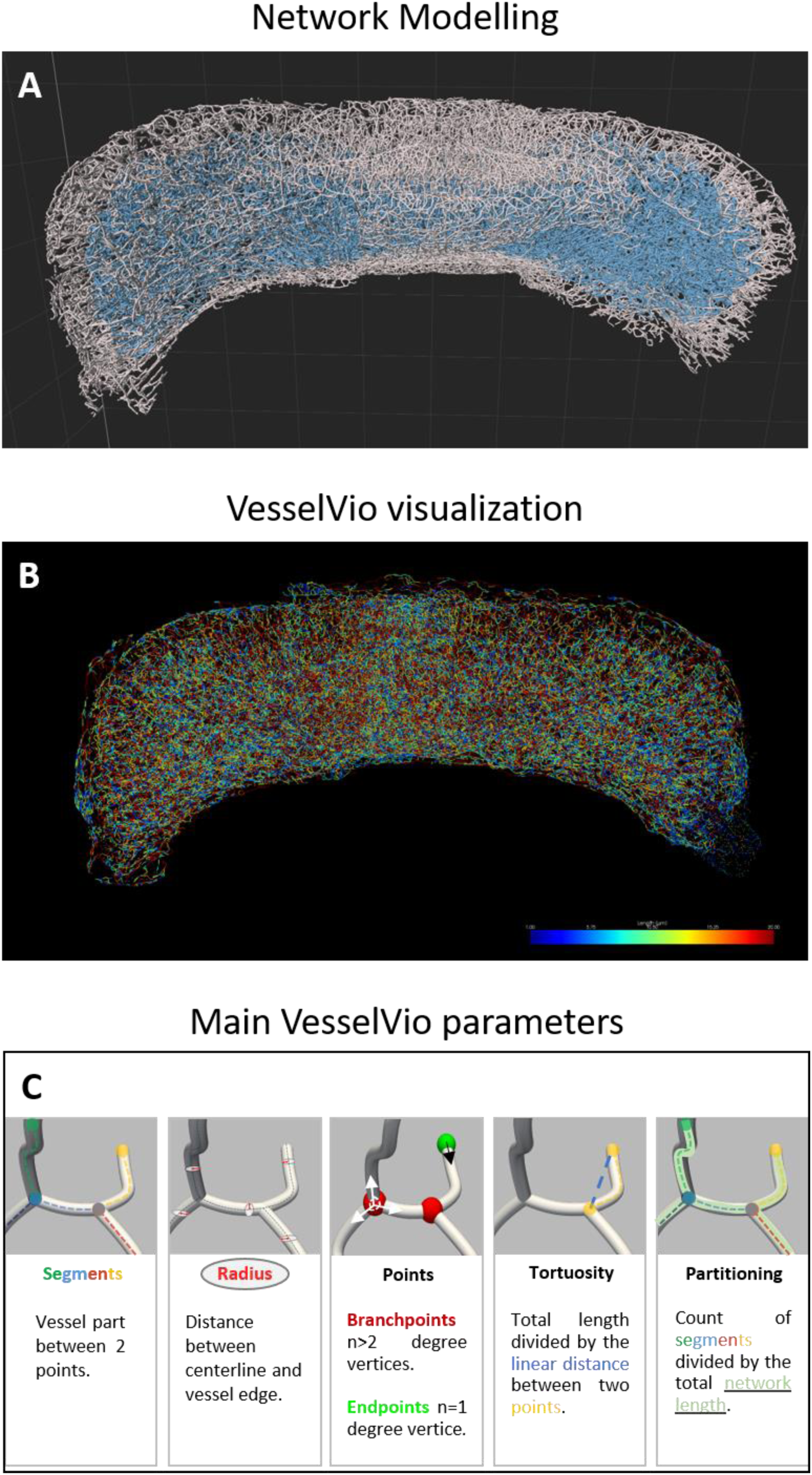
Illustration of the vascular network modeling of a postnatal mouse cerebellum and the parameters defined on VesselVio software. A,B : Illustration of the matching between the Imaris network modelling (A) and the corresponding VesselVio visualization (B). In A, the superficial network is represented in white and the deep network is represented in blue. In B, the different ranges of segment length are indicated by the colored scale. C: Graphical definition of the network parameters analyzed with VesselVio.

Then, for each network (deep and superficial), a channel was created from the filament modelling via a Matlab macro linked to Imaris (*Image Processing → create channel from Filaments*) and then opened in the Fiji software via the *Image to Fiji* bridge. Thus, the files were converted in 8-bit images, binarized, and saved into *.tiff* and *.nii* formats. Each file is then opened in the VesselVio software and the vascular skeletons were visualized in .*tiff* files to check file conversion before analyzing (Figure 3B). Then, all segmented vasculature datasets were downloaded in .*nii* files for analysis. Finally, a single excel file is created with the specific parameters of each vasculature. In this study, the parameters of interest were i) the total volume, area, length and number of segments, for the whole cerebellar network, ii) the mean volume, area, length and radius, for segments, and iii) some specific vessel characteristics such as the branchpoints, endpoints, tortuosity and segment partitioning (Figure 3C).

### Effect of IH on the expression of angiogenesis-related factors

The transcriptomic analysis was targeted at genes known to be involved in angiogenesis and performed on whole cerebella at different stages of postnatal development (Figure 4). We tested a panel of 23 genes, of which 22 were differentially regulated in at least one stage. The exception being Col1a1 which did not vary across our experiments, as such, it is not represented in Figure 4. Overall, we found no sex- or litter-dependent effect.

**Figure 4.**
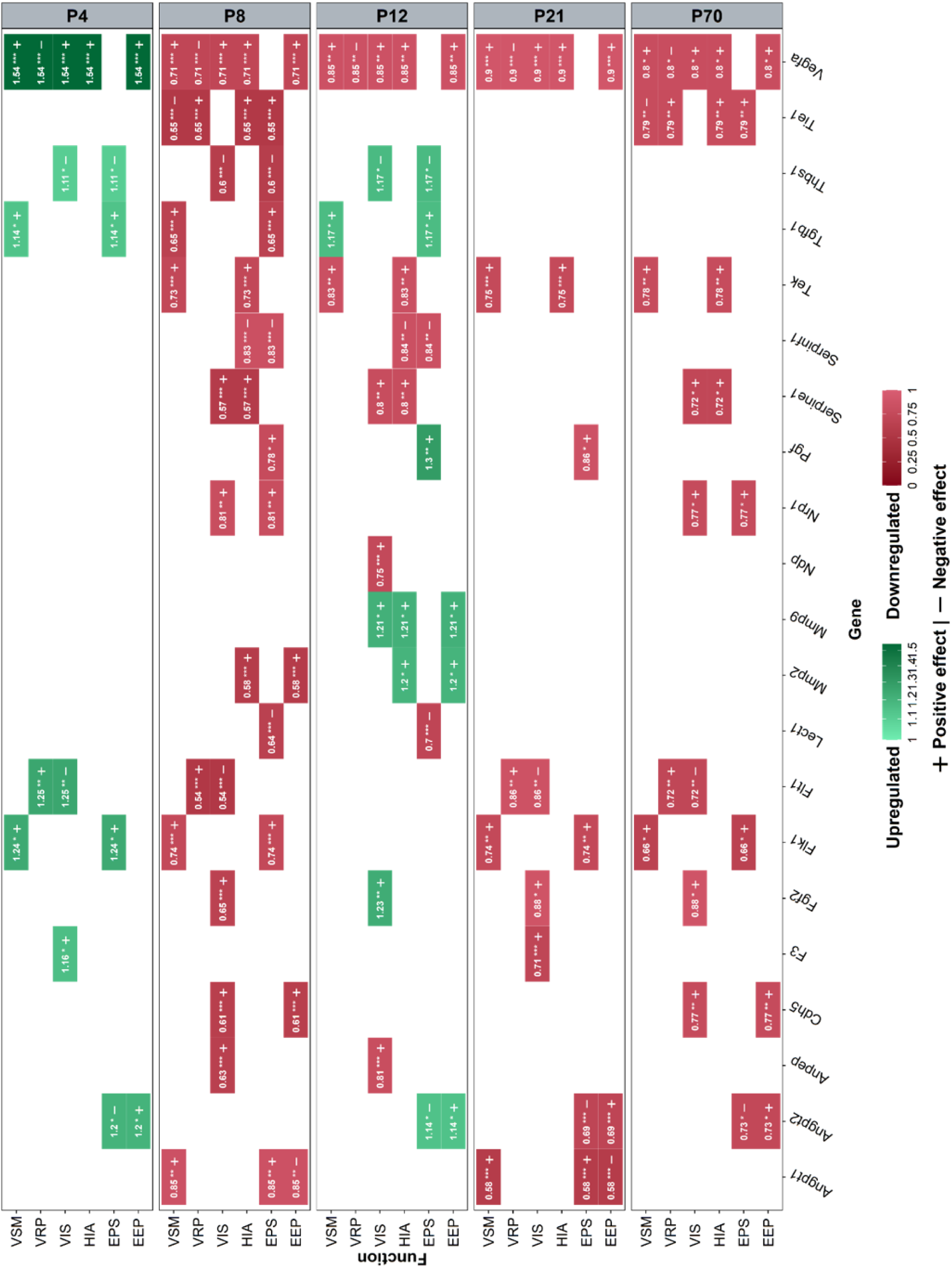
(next page): Effect of intermittent hypoxia on angiogenesis-related genes during cerebellar development. Significant results reflecting the regulation of the expression of genes involved in cerebellar angiogenesis after IH in P4, P8, P12, P21, and P70 mice (right y-axis) measured by qRT-PCR. Upregulated (green) and downregulated (red) genes are represented for each pathway and the color corresponds to the gradient of the fold change value indicated in each box. Each gene (x-axis) can be mapped back to one or several functional pathways (left y-axis). EEP: extracellular matrix and endothelial permeability; EPS: endothelial proliferation and survival; HIA: hypoxia-induced angiogenesis; Px: postnatal day x; VIS: vascular invasion and sprouting; VRP: vascular remodeling and patterning; VSM: vascular stabilization and maturation. Asterisks indicate the level of statistical significance: one for p < 0.05, two for p < 0.01, and three for p < 0.001.

At stage P4, 30.4% of the genes were regulated (7/23 genes), all of which are upregulated in the IH condition. Most notably, Vegfa and two VEGF receptors Flk1 and Flt1 are overexpressed, indicating the importance of this pro-angiogenic signaling pathway at this stage. Interestingly, some negative regulators of angiogenesis are also overexpressed, such as endothelial adhesion molecule thrombospondin 1 (Thsb1).

P8 is the most regulated stage overall with 73.9% (17/23) differentially expressed genes, but in contrast with P4, all were downregulated under IH. This includes the previously P4 upregulated growth factors but also nearly all genes we tested from the angiopoietin and VEGF signaling pathways. In parallel to this effect on pro-angiogenic molecules, IH also acts negatively on some anti-angiogenic factors such as Thbs1 and chondromodulin1 (Lect1). Moreover, despite being 6 days into the hypoxia protocol, genes involved in hypoxia induced angiogenesis, such as Vegfa and the angiopoietin 1 inhibitor Tie1, are also downregulated at P8.

Then, P12 had 60.9% (14/23) of regulated genes with a partial switch to upregulation compared to P8. This is the case of the growth factors Fgf2, Pgf and Tgfb1 but also the anti-angiogenic Thsb1. Interestingly, the results showed that IH upregulates the expression of matrix metalloproteases 2 and 9 involved in vessel invasion and sprouting.

The decrease in IH-regulated genes continues at P21 with only 39.1% (9/23) differentially expressed genes. All of them are under-expressed, including Flt1, a negative regulator of neovascularization. Surprisingly, 43.5% (10/23) were still regulated at P70, suggesting that the vascular response to hypoxia is long lasting on a transcriptomic level. This long-term effect decreases angiogenic mechanisms, especially with the downregulation of hypoxia-induced angiogenic factors and positive regulators of vascular invasion and sprouting.

### Effect of IH on the vascularization of the postnatal cerebellum

Because of the significant inter-individual variability, all the following results were normalized relative to the total volume of the cerebellum.

Concerning the general parameters of the entire vascular network, the IH protocol induced some major modifications at P4 (Figures 5 and 6), with a strong increase in the volume, area and length observed at P4 in IH condition, indicating a denser network from the very first days (Figure 5). The higher number of segments, branchpoints and partitioning confirms the increased complexity of the vascularization after hypoxia. In contrast, the tortuosity is decreased at P4 in hypoxic animals, indicating an inhibiting effect on vessel remodeling (Figure 6). Thanks to our analysis workflow (Figure 2F), that makes a differential analysis of the superficial and deep networks possible, we showed that the differences observed for the volume, the length and the partitioning are mainly due to an effect of IH on the superficial network, whereas the increase of the segments and branchpoints is attributable to the deep vessels (Figures 8-9). From P8 onwards, all the general IH-induced modifications are attenuated and become mostly non-significant (Figures 5-6), suggesting that hypoxia no longer has an effect on the cerebellar vasculature at later stages. However, our differential study revealed a specific increase of the deep vessel partitioning at P8, and of the length, area and partitioning of the superficial network at P12 (Figure 8-9). At P21, only the number of endpoints is affected by IH (Figure 6) but, by looking at the two networks separately, it appears that this decrease is also present at P8 and mainly affects the deep vessels (Figure 9).

**Figure 5:**
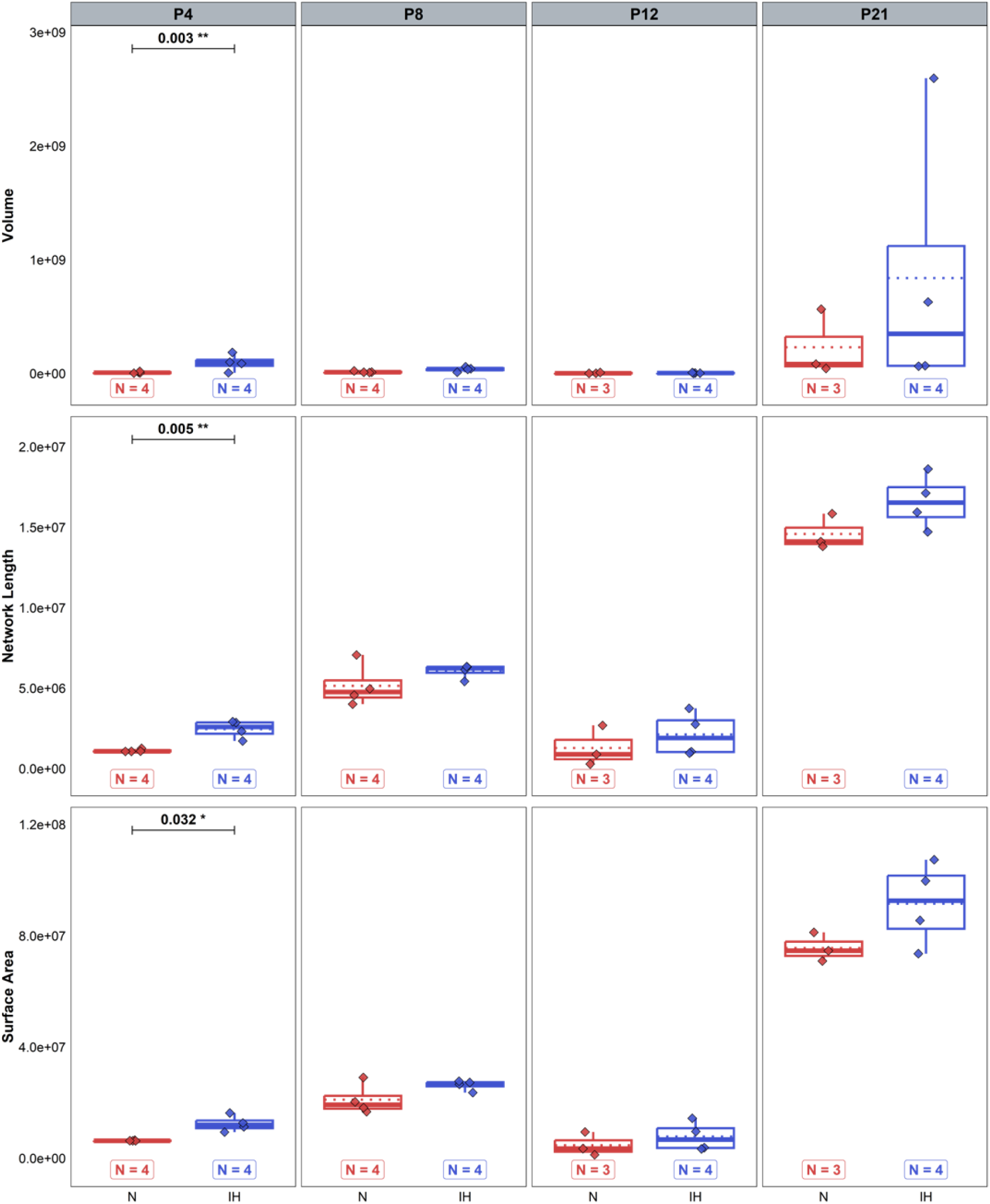
Effect of intermittent hypoxia on global vascular parameters of the cerebellum across stages. Measurement of the volume, length, and surface area of the cerebellar global vascular network in control (red) and hypoxic (blue) mice at P4, P8, P12, and P21. For each variable, the total number of animals in each experimental group is indicated under the boxplots and represented by diamonds. Exact p-values are indicated above the plot when significant. IH: intermittent hypoxia; N: normoxia; Px: postnatal day x.

**Figure 6:**
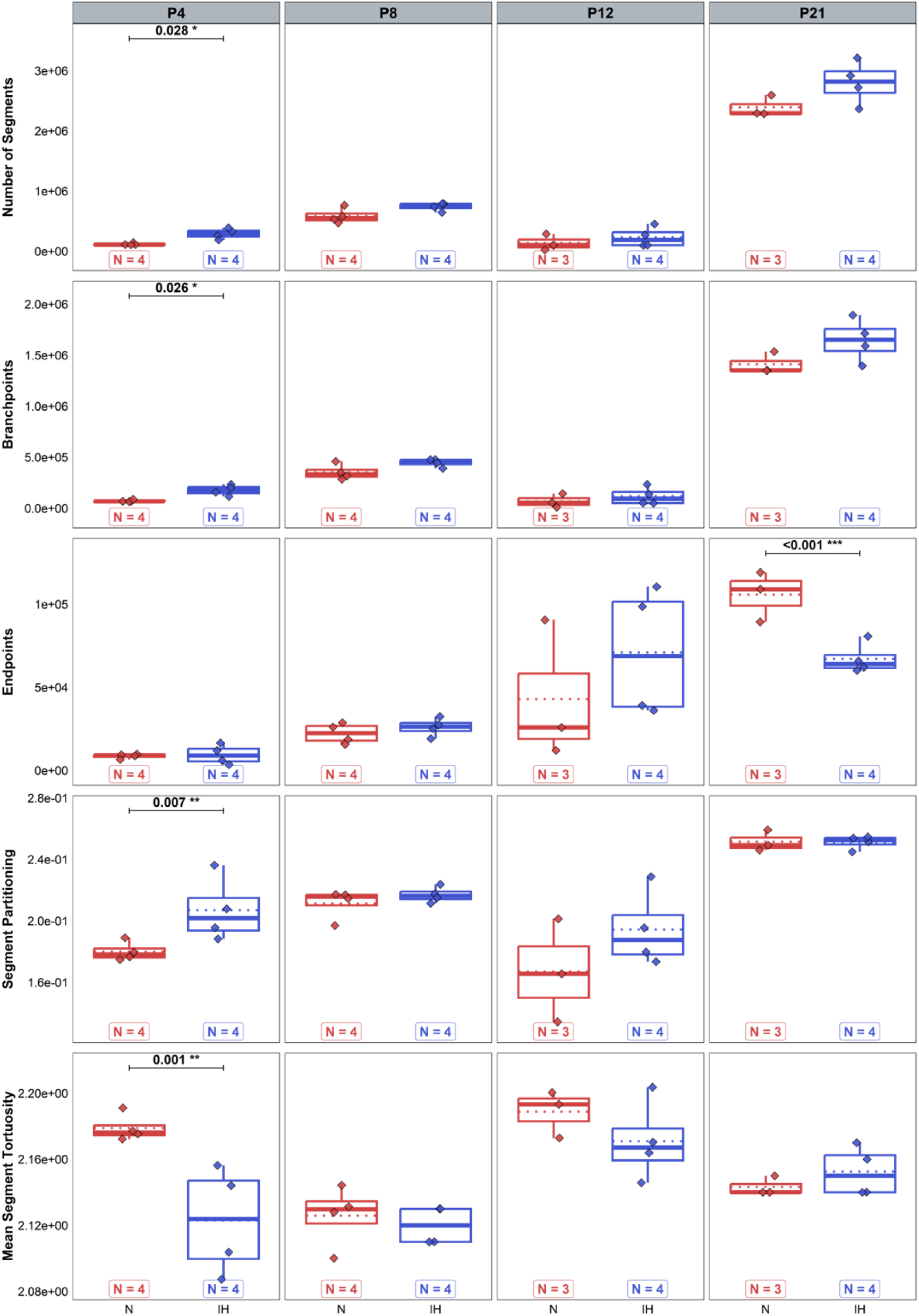
Effect of intermittent hypoxia on vascular segment parameters of the cerebellum across stages. Measurement of the number of segments, branchpoints, endpoints, segment partitioning, and tortuosity in the overall cerebellar vascular network in control (red) and hypoxic (blue) mice at P4, P8, P12, and P21. For each variable, the total number of animals in each experimental group is indicated under the boxplots and represented by diamonds. Exact p-values are indicated above the plot when differences are significant. IH: intermittent hypoxia; N: normoxia; Px: postnatal day x.

Meanwhile, the effects of IH on segment characteristics are more persistent and variable between ages. At P4, all mean segment parameters are affected by IH with an increase of the mean volume but a concomitant decrease of the area, length and radius (Figure 7). All these variations are mainly due to the superficial network, whose tortuosity is also specifically altered (Figure 9). At P8, we only observed a continuing higher mean volume of segments after IH in studying both networks together (Figure 7) but the differential analysis showed a decreased mean length of deep segments and an increased mean area of superficial segments, which are cancelled by a non-significant opposite effect of IH on the deep network (Figure 10). At P12, hypoxia induced a decrease of the mean segment volume, area and length largely attributable to the superficial vessels (Figures 7-10). Finally, at P21, only slight alterations remained with a higher tortuosity of the superficial vascular segments (Figure 9).

**Figure 7:**
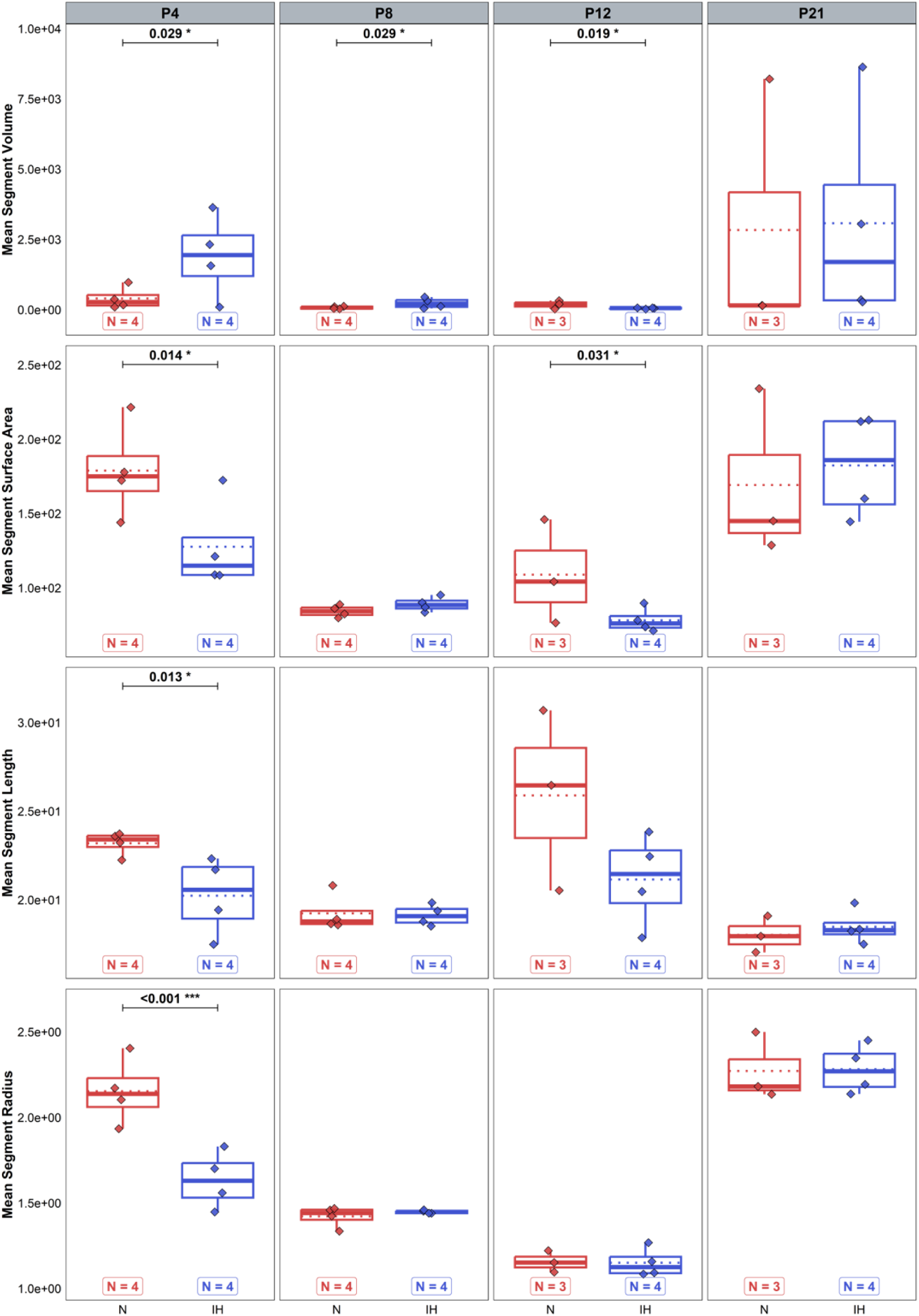
Effect of intermittent hypoxia on mean segment parameters of the cerebellum across stages. Measurement of the mean volume, surface area, length, and radius of segments in the overall cerebellar vascular network in control (red) and hypoxic (blue) mice at P4, P8, P12, and P21. For each variable, the total number of animals in each experimental group is indicated under the boxplots and represented by diamonds. Exact p-values are indicated above the plot when differences are significant. IH: intermittent hypoxia; N: normoxia; Px: postnatal day x.

**Figure 8:**
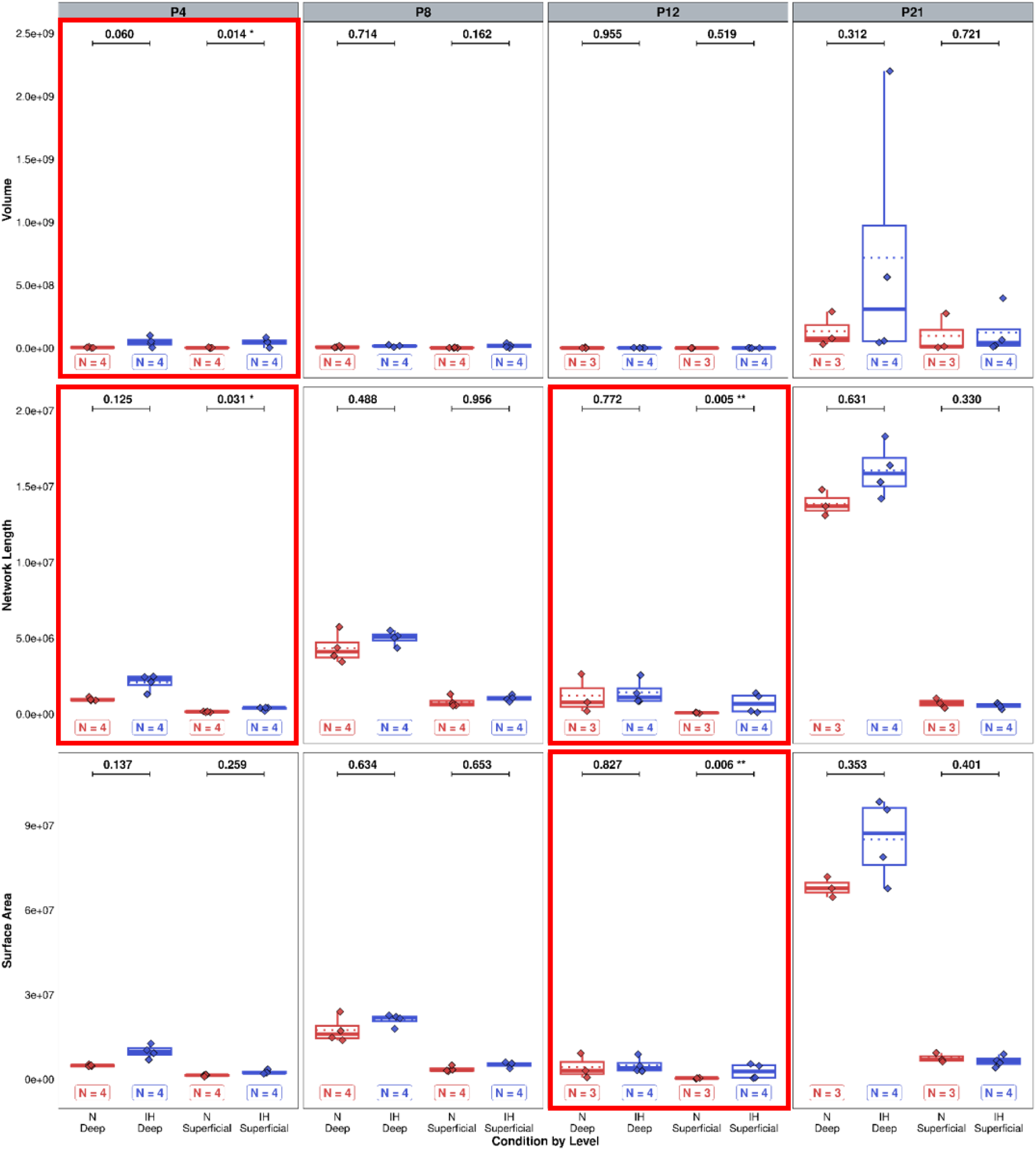
Effect of intermittent hypoxia on global parameters of the superficial and deep vascular networks of the cerebellum across stages. Measurement of the volume, length, and surface area of the cerebellar vascular superficial and deep networks in control (red) and hypoxic (blue) mice at P4, P8, P12, and P21. Box-plots framed in red show the differential effect of hypoxia on the two networks. For each variable, the total number of animals in each experimental group is indicated under the boxplots and represented by diamonds. Exact p-values are indicated above the plot. IH: intermittent hypoxia; N: normoxia; Px: postnatal day x.

**Figure 9:**
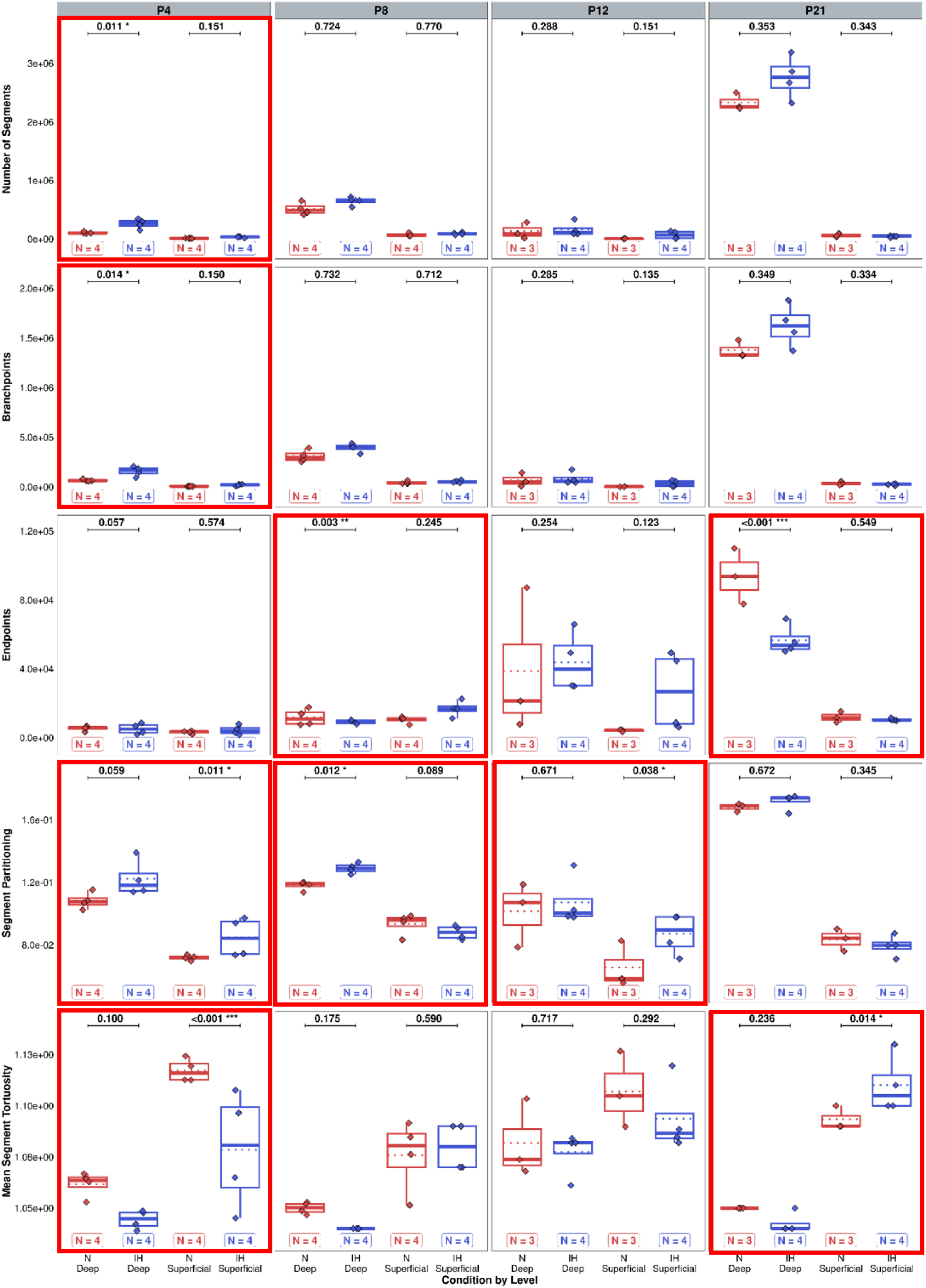
Effect of intermittent hypoxia on segment parameters of the superficial and deep vascular networks of the cerebellum across stages. Measurement of the number of segments, branchpoints, endpoints, segment partitioning, and tortuosity of the cerebellar vascular superficial and deep networks in control (red) and hypoxic (blue) mice at P4, P8, P12, and P21. Box-plots framed in red show the differential effect of hypoxia on the two networks. For each variable, the total number of animals in each experimental group is indicated under the boxplots and represented by diamonds. Exact p-values are indicated above the plot. IH: intermittent hypoxia; N: normoxia; Px: postnatal day x.

**Figure 10:**
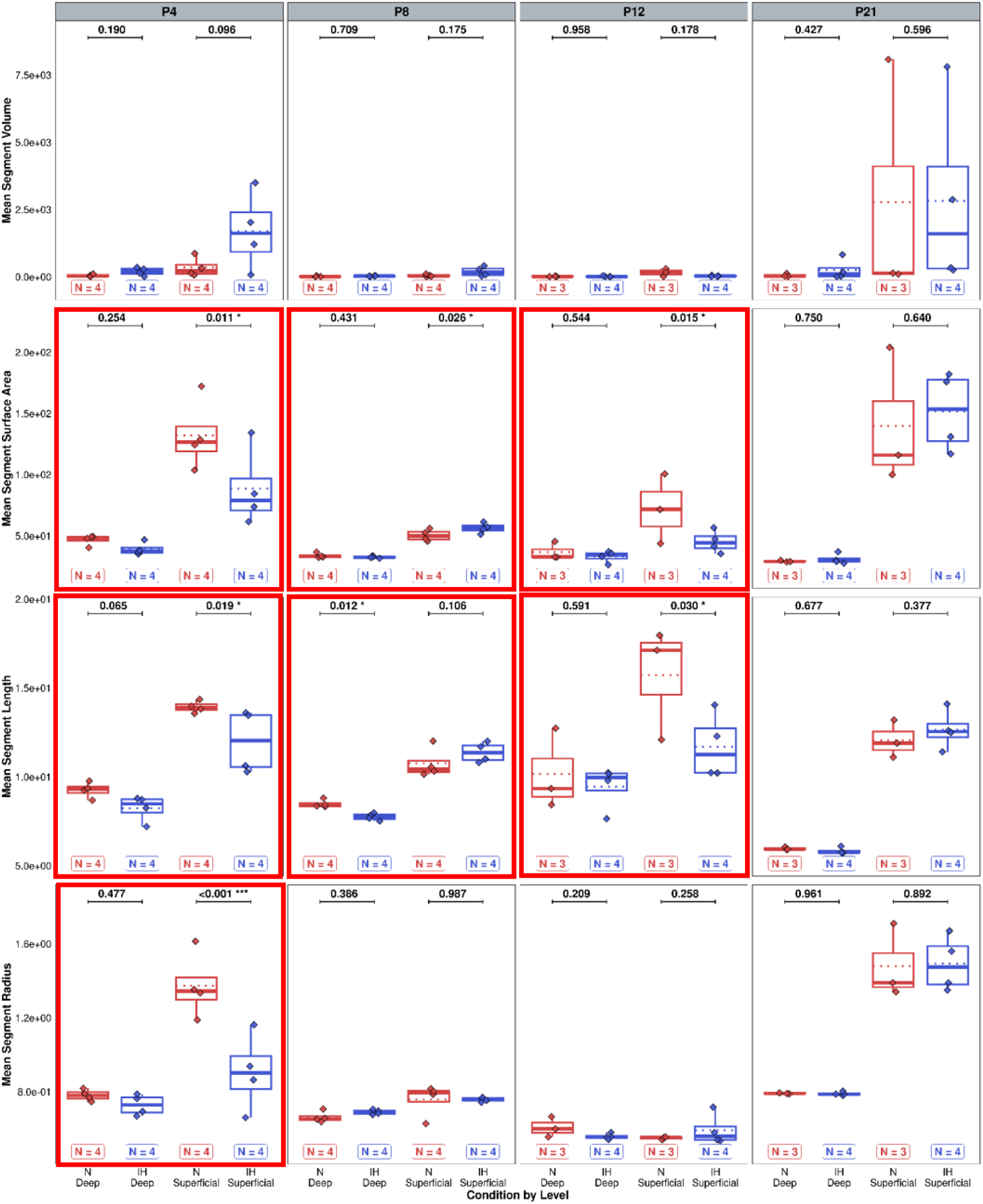
Effect of intermittent hypoxia on mean segment parameters of the superficial and deep vascular networks of the cerebellum across stages. Measurement of the mean segment volume, surface area, length, and radius of the cerebellar vascular superficial and deep networks in control (red) and hypoxic (blue) mice at P4, P8, P12, and P21. Box-plots framed in red show the differential effect of hypoxia on the two networks. For each variable, the total number of animals in each experimental group is indicated under the boxplots and represented by diamonds. Exact p-values are indicated above the plot when significant. IH: intermittent hypoxia; N: normoxia; Px: postnatal day x.

## Discussion

### An optimized method to visualize the cerebellar vascular network in 3D

Despite the advancements in the study of cerebellar neurogenesis, we still lack knowledge regarding its angiogenesis during development. This lack of information is mainly due to the difficulties in accessing the cerebellar depths with conventional imaging techniques. Fortunately, the advancement of clearing protocols and light sheet microscopy in the past decade has gradually enabled the 3D visualization of the vasculature of the cerebellum in its earliest stages as well as in later ages (Renier at al., 2014; Ueda et al., 2020). However, the analysis of these 3D images is still difficult because only a few software programs can model and produce quantitative data from such 3D files.

One of the typically used software packages for 3D modelling is IMARIS from Oxford Instruments Company. However, due to the complexity and the extreme variability of cerebellar vasculature, the IA module is not efficient to model the whole network at once. Therefore, a threshold of intensity needs to first be applied to separate vessels into a superficial network containing the large vessels (>20-µm diameter) and a deep network consisting of the small deep capillaries. Then machine learning can be used to optimize modelling, but these learning steps must be limited because errors quickly increase due to the accumulation of conflicting information. Furthermore, although this software is the most efficient for modelling, its statistical tools are not adapted to the study of vascular networks. It is therefore necessary to turn to other software solutions to obtain quantitative data. Vesselvio (Bumgarner and Nelson, 2022) proved to be a good solution, as each vascular parameter is well defined. Furthermore the software is open access and has already been used in other vascular studies. Therefore, after numerous steps of optimization and software tests, a practical, optimal, and user-friendly workflow was created for 3D vascularization analysis.

### Early changes

Our innovative 3D image analysis approach allowed the assessment of the evolution of several vascular parameters in response to our hypoxia protocol. The most changes in the cerebellar vascular network features appear at the P4 stage, after only two days of hypoxia. Overall, the network of IH mice has a higher volume, length and surface area than the control animals mostly attributable to the superficial network. This increase is consistent with the upregulation of several factors promoting vessel growth such as Vefga (Molina et al., 2013) and its VEGF receptor Flk1 (Huang et al., 2015; Wang et al., 2014), transforming growth factor beta 1 (Tgfb1; Molina et al., 2013) or F3 (Peña et al., 2017). This suggests that an acute response to hypoxia at this early stage involves grossly increasing vascularization to compensate for IH exposure. In contrast, the decreased tortuosity of vessels in IH mice, especially in the superficial network, could suggest less remodeling. This hypothesis can be supported by the overexpression of two negative regulators of vascular invasion and sprouting (VIS), namely Flt1 (Nesmith et al., 2017; Wild et al., 2017) and Thbs1 (Lawler and Lawler, 2012).

The P4 enlarged network is composed of more numerous segments with more branchpoints, especially in the deep network, accompanied by an increased segment partitioning, predominantly in the superficial network. However, the vascular segments of IH mice are shorter and thinner, predominantly in the superficial network. This observation supports the early emphasis on increasing the network rather than refining it after hypoxia, and suggests that hypoxia-induced angiogenesis (HIA) leads to a more complex but less mature vascular network. This hypothesis also correlates with the high upregulation of Vegfa which is a known actor of HIA (Nesmith et al., 2017) as well as of Angpt2 which promotes EEP in response to hypoxia (Trollmann et al., 2018; Vates et al., 2005) and of F3 which stimulates neovascularization (Heidari et al., 2024; Peña et al., 2017).

### Vascular adaptation during IH

In contrast to P4, no significant difference in general network parameters was observed at P8, and only the increase in the mean segment volume is maintained after 4 days of IH. In parallel, all tested genes were downregulated in all functional groups, including the previously upregulated growth factors, but also a large panel of positive regulators of angiogenesis. Among them, we can notice the strong under-expression of the angiogenic receptor Tek and its regulator Tie1 (Lee et al., 2009; Savant et al., 2015), of metalloprotease 2 (MMP2; Quintero-Fabián et al., 2019; Sang, 1998), of alanyl aminopeptidase (Anpep; Rangel et al., 2007) and of Serpine 1 (Zhang et al., 2023), which are all known to be implicated in HIA. This result suggests that the initial acute angiogenesis triggered at the beginning of the protocol in response to IH has ceased by P8. In accordance with this hypothesis, our differential study across networks reveals that the length and the surface area of the superficial vessels increase with a slight decrease of the partitioning. This indicates that the network is still growing but is less complex with fewer vessel interconnections. In contrast, a remodelling process is ongoing in the depth of the cerebellum as an increase of the vessel partitioning associated with a decrease of the number of endpoints and of the mean segment length are observed. This could be linked to the under-expression of some negative regulators of VIS, namely Flt1 and Thbs1 (Lawler and Lawler, 2012; Nesmith et al., 2017; Wild et al., 2017) or of endothelial cell mechanisms such as Serpinf1 (He et al., 2015; Tombran-Tink, 2005) or Angpt1 (angiopoietin 1; Lee et al., 2009; Zacharek et al., 2007).

At P12, like at P8, general parameters are not altered but a decrease of the mean segment volume and area is observed, suggesting that the overall shape of blood vessels stabilizes during the protocol. However, while the superficial vascular network was only slightly affected at P8, it shows several hypoxia-related alterations at P12, namely an increase in length, area and partitioning. This complexification of the network could be explained by a partial switch in the expression of several genes. Indeed, while some pro-angiogenic factors remain under-expressed, others become over-expressed such as the matrix metalloproteases MMP2, MMP9, and Angpt2, which make the extracellular matrix more permissive to vessel endothelial tip cells (Cai et al., 2017; Quintero-Fabián et al., 2019; Sang, 1998; Vates et al., 2005). In addition, norrin (Ndp), which was previously not affected by IH, becomes under-expressed, magnifying the effect on the matrix (Luhmann et al., 2008; Ohlmann et al., 2010; Ohlmann and Tamm, 2012; Zhang et al., 2017). Taken together with the upregulation of Fgf2, Pgf and Tgfb1, involved in EPS and VSM, this reveals the emergence of a second phase of IH response at later stages of development (Lee et al., 2023; Luna et al., 2016; Molina et al., 2013).

Ten days after the end of the IH protocol, P21 IH mice present a lower number of vessel endpoints than their normoxic counterparts, largely attributable to the deep network. This coincides with the emergence of newly downregulated VIS factors such as the critical Angiopoietin 2 (Angpt2; Raybaud, 2010) but also F3 (Heidari et al., 2024; Peña et al., 2017), Fgf2 (Mori et al., 2017), as well as Cdh5 (Bentley et al., 2014; Nan et al., 2023; Wallez et al., 2006) at P70. At P21, the cerebellum is also characterized by a higher tortuosity of superficial vascular segments, indicating that, despite compensatory mechanisms occurring at the end of the IH protocol, minor alterations to the vascular network remain visible in the long term. According to this observation, the majority of genes under-expressed at P21 are still under-expressed at P70, including Flk1, Tek, and Vegfa (Nesmith et al., 2017; Zacharek et al., 2007).

### Vasculogenesis and cerebellar development during IH

The brain, being so energy demanding, is particularly dependent on vascular function, thus its growth is hindered by poor blood supply (Donkelaar, 2014). Our results confirm that it is also the case for the cerebellum, as they demonstrate that IH induces an alteration of the cerebellar angiogenesis associated with a delay in cerebellar maturation (Leroux et al., 2022). Moreover, beyond the classical role of the vasculature as O_2_ supplier, it is now well known that the crosstalk between blood vessels and nerve cells is also critical for various cell processes which occur during brain development such as migration, differentiation or precursor proliferation (Segarra et al., 2019). As cerebellar development takes place mainly during the postnatal period in both mice and humans, the vascular abnormalities induced by our IH protocol could lead to the cellular disorganization observed at P12 in mice that underwent hypoxia (Leroux et al., 2022).

Moreover, neurons and endothelial cells are reported to share a large array of cellular pathways (Larrivée et al., 2009; Weinstein, 2005). Thus we cannot exclude that both neural and vascular systems are affected in parallel by IH. For example, NRP1 is known to be a guidance cue for both neurons and blood vessels (Fantin et al., 2009; Vieira et al., 2007) by either binding VEGFA (Erskine et al., 2011) or SEMA3A (Eichmann et al., 2005) and the expression of these three factors are significantly decreased during our IH protocol (Rodriguez-Duboc et al., 2023). We can also cite the matrix metalloproteases, which are involved, not only in brain angiogenesis (Girolamo et al., 2004) but also in the morphogenesis of the cerebellar cortex (Luo, 2005).

From a cellular standpoint, the parallels between vascularization and neurite development also confirm our previous findings on Purkinje cell arborization. Indeed, the Angpt2/Tie1 pathway was highly regulated by IH. This system is well known to participate in vessel formation (Augustin et al., 2009) but has been more recently described as a regulator of dendritic tree development in Purkinje cells (Luck et al., 2021). Given our findings on the alterations of Purkinje cell morphogenesis after IH (Leroux et al., 2022; Rodriguez-Duboc et al., 2023), this cements the idea of a parallel alteration of neurogenesis and angiogenesis during IH. Finally, despite few observable differences in the vascular network of IH mice at P21, the expression of numerous genes implicated in this system is still dysregulated and may contribute to behavioral, histological and neuronal deficits we have shown in adults after our IH protocol (Leroux et al., 2022; Rodriguez-Duboc et al., 2023).

## Conclusion

Taken together with our previous studies, this work confirms that neurogenesis and angiogenesis are extremely interlocked during cerebellar postnatal development as during brain embryogenesis. Moreover, both systems are affected by a perinatal IH with the alteration of both vascular network shape and cerebellar cortex histogenesis. Thus, our results indicate that, in pathologies such as apnea of prematurity, mimicked by the present protocol (Cai et al., 2012), the two components participate in short and long-term deficits observed in patients. More generally, when treating patients with neurodevelopmental disorders, it would be a mistake to focus therapeutic approaches solely on neurological damage, without taking into account the vascular aspect.

## Declarations

### Availability of data and code

The raw datasets, analyses and code supporting this work can be accessed via the GitHub repository available at https://github.com/agalic-rd/Vasc-AoP, also referenced on Zenodo under: https://doi.org/10.5281/zenodo.15319299.

### CRediT author statement

**Agalic Rodriguez-Duboc:** Investigation, Methodology, Project administration, Data Curation, Formal analysis, Visualization, Validation, Writing - Original Draft, Writing - Review and Editing. **Camille Racine:** Investigation, Writing - Original Draft, Writing - Review and Editing. **Magali Basille-Dugay:** Supervision, Resources, Writing - Review and Editing. **David Vaudry:** Funding acquisition, Resources, Writing - Review and Editing. **Bruno Gonzalez:** Funding acquisition, Resources, Writing - Review and Editing. **Delphine Burel:** Investigation, Conceptualization, Methodology, Funding acquisition, Project administration, Supervision, Resources, Writing - Original Draft, Writing - Review and Editing.

## Acknowledgments

We acknowledge the France-BioImaging infrastructure (https://ror.org/01y7vt929) supported by the French National Research Agency (ANR-24-INBS-0005 FBI BIOGEN). We are also grateful to François Chadelaud and Dr. Sarah Leroux from Rouen University for developing the hypoxia chamber which is the basis of this work. Finally, we would like to thank Dr. David Godefroy for his help with the 3D image acquisition and Alexis Lebon for his help with the software.

## Supplementary data

**Supplementary Table 1:**
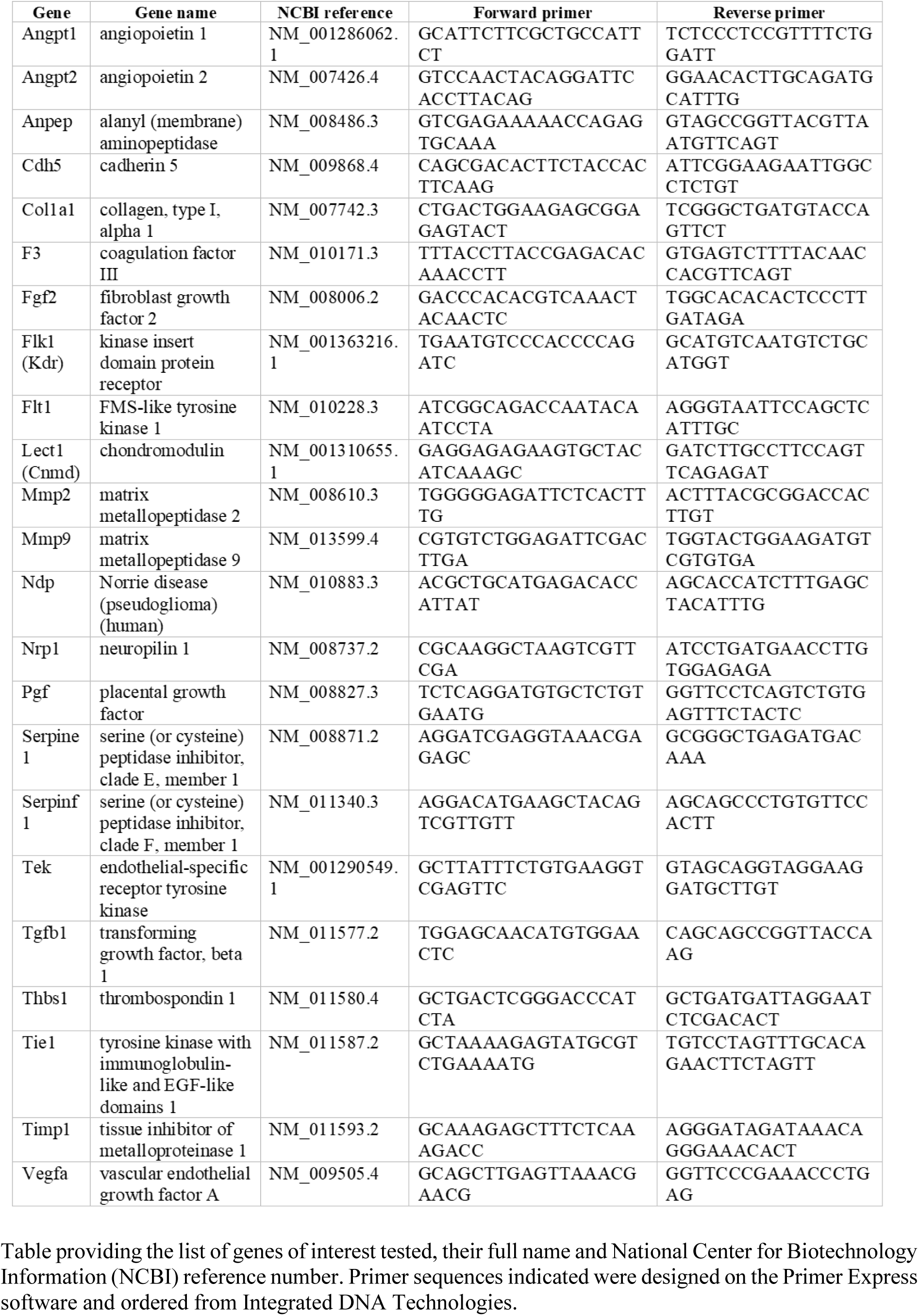
List of primers used for qRT-PCR experiments and their corresponding sequences.

**Supplementary Table 2.**
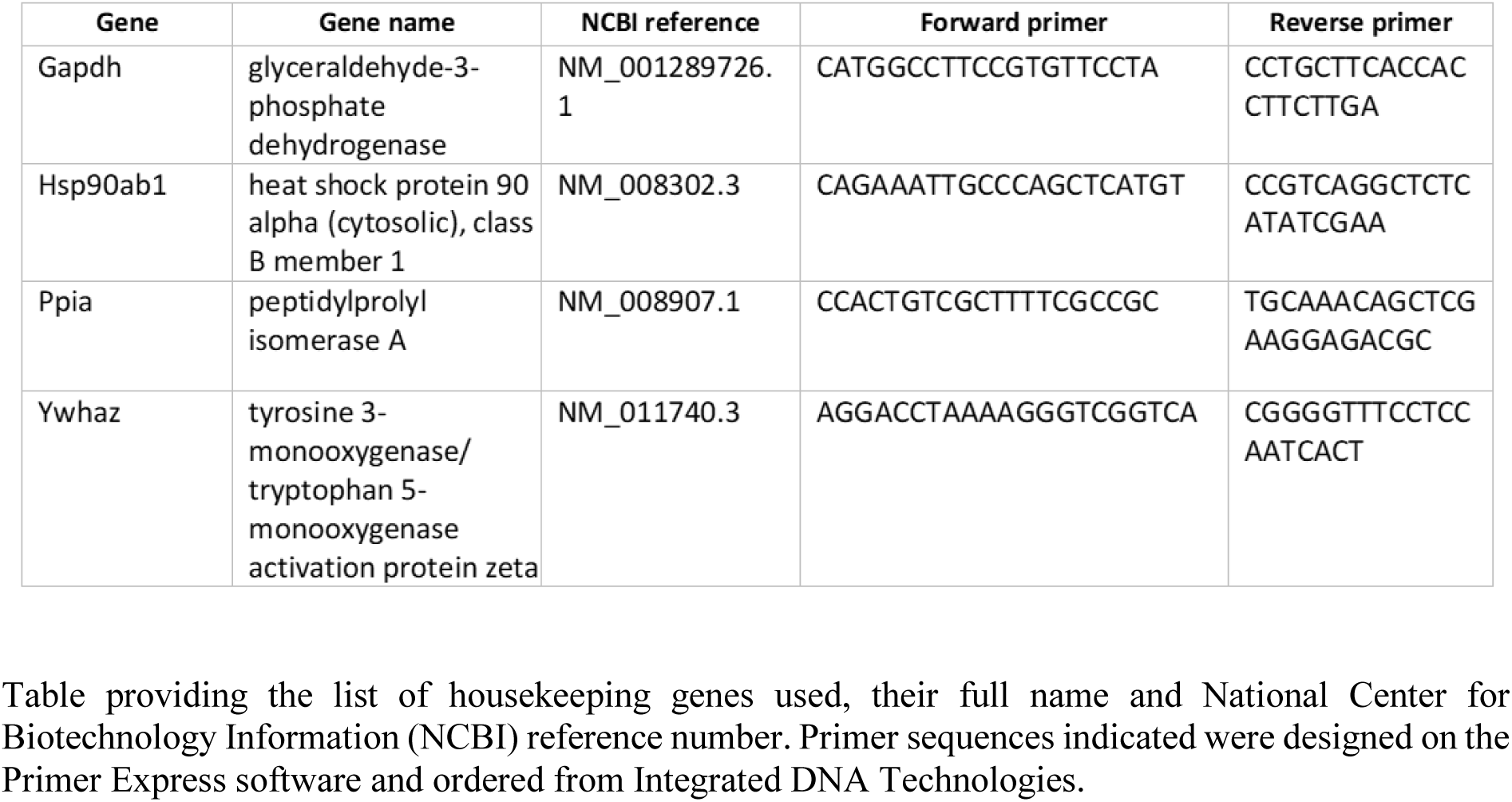
Selected housekeeping genes for qRT-PCR experiments.

